# Time mating guinea pigs by monitoring changes to the vaginal membrane throughout the estrus cycle and with ultrasound confirmation

**DOI:** 10.1101/2020.08.21.256255

**Authors:** Rebecca L Wilson, Kristin Lampe, Brad J Matushewski, Timothy RH Regnault, Helen N Jones

**Author notes:** Corresponding Author: Rebecca Wilson, College of Medicine, Department of Physiology and Functional Genomics, University of Florida, Gainesville, Florida 32610.

## Abstract

Guinea pig development *in utero* is more similar to humans than any other rodent species. As such, their importance to reproductive studies is evident, particularly those studies focused on therapeutic interventions to improve human pregnancy outcome. Tracking the guinea pig estrus cycle is imperative to ensuring appropriately timed mating and can be performed by monitoring the guinea pig vaginal membrane. Here, we describe, and provide picture representation, of changes to the guinea pig vaginal membrane throughout the estrus cycle. Utilization of this monitoring enabled a 100% pregnancy success rate on the first mating attempt in a cohort of five guinea pigs. This approach, along with early pregnancy ultrasounds as a secondary method to confirm pregnancy, offers a cost effective and reliable approach to timed-mating in the guinea pig.

## Introduction

The placenta is a transient organ, unique to pregnancy which functions to support the passage of oxygen and nutrients to the developing fetus and maintain adaptation of the maternal environment to pregnancy [1]; without it, there is no pregnancy. Inappropriate placental development and function is associated with numerous obstetric complications which affect 1 in 4 pregnancies worldwide [2]. Conditions such as preeclampsia, stillbirth and preterm birth contribute significantly to both maternal and neonatal morbidity and mortality. As such, there is a heightened need to continue to develop robust predictive models, diagnostic techniques, and potential therapies for use during pregnancy. However, studying and utilizing pregnant women is considered high risk and consequently, most pregnancy research utilizes animal models from small rodent species to larger mammals such as sheep.

The usefulness of the guinea pig in pregnancy research is often overlooked and more readily performed in mice due to their shorter gestation, lower costs and more rapid advancement of transgenic model development. However, guinea pigs have smaller litters and deliver precocial young (they are born fully-furred with well-developed sensory and locomotory abilities) [3]. Further still, guinea pigs have some similar developmental milestones to humans both throughout gestation and following birth [3, 4]. This includes significant fetal fat deposition in late pregnancy [5] and comparable biochemical and morphological development of fetal organs including lungs [6], kidneys [7] and the cardiovascular system [8]. Unlike rats and mice, the type of placenta in the guinea pig reflects the human as they have a haemomonochorial placental barrier: a single syncytiotrophoblast layer separates the maternal blood space from the fetal vessels [9]. Placental establishment is also more similar to humans, with guinea pigs exhibiting deeper placental trophoblast invasion then mice [10] and changes to the maternal hormonal profile throughout pregnancy more closely resembles changes in humans when compared to other rodent species [11]. Altogether, these outcomes make utilizing the guinea pig in pregnancy research superior, particularly for the advancement of understanding the potential mechanisms underlying poor obstetrical outcomes. This knowledge can then be used to facilitate the development of potential therapies to treat obstetrical complications that significantly contribute to long-term morbidity and mortality for mother and child, and ultimately reduce the impact on metabolic health in later life.

One of the principal challenges associated with using guinea pigs for reproductive discoveries is the ability to time their matings. Unlike other rodent species, the formation of a copulatory plug by male guinea pigs after intercourse is debated [12] and as such, it is difficult to ascertain when mating has occurred in order to track pregnancy progression. Some have described methods in which time-mating guinea pigs is achieved by allowing females to carry a pregnancy and then breed in 24 h immediately after birth [13]. Others simply allow a time range of plus/minus 1 week based on the length at which the female is housed with the male [14]. However, these methodologies introduce significant variability (mean fetal weight increases from 2.5 g at GD33 to 12.0 g at GD39; R.Wilson unpublished) and time which ultimately increases costs associated with performing experiments with guinea pigs. Like other rodents, guinea pigs have a four-stage estrus cycle: estrus, metestrus, diestrus and proestrus [15]. Estrogen is the dominant hormone during the estrus period and is when ovulation occurs, whilst progesterone dominates the diestrus period [16]. All stages of the estrus cycle can be monitored through vaginal cytological smears however, these can be time consuming and require experience in understanding the technique. Vaginal impedance can also be used to monitor estrus cycles progression [17] but is also another technique that requires specialized tools. Herein, we describe a simple method to achieve timed-mated guinea pig pregnancies based on monitoring changes in the vaginal membrane (which perforates spontaneously at estrus) [12], and ultrasound in early pregnancy (gestational day 21-30).

## Results

### Guinea pigs undergo a ∼16 day cycle in which the vaginal membrane perforates during time of ovulation

We took 5 female Hartley guinea pigs between 500-550 g (8-10 weeks of age) and began daily monitoring of the estrus cycle by observing changes to the vaginal membrane. During the latent period of the estrus cycle, the vaginal membrane was a pale color and visibly closed (Figure 1A & B). In the 4-5 days prior to ovulation, the vagina underwent noticeable changes in color; becoming progressively darker pink; mild swelling of the vulva could also be noted prior to perforation (Figure 1C). Full perforation of the vagina happened within 24-72 h and was noticeable as a complete opening of the vagina, often associated with increased vaginal secretions/mucous (Figure 1D). The vaginal membrane then closed within 48-72 h of full perforation (Figure 1E & F). Based on these observations, we confirmed the guinea pig estrus cycle to be 15-16 days (n=4 out of 5 female guinea pigs) with observations of initial perforation through to closing of the vaginal membrane lasting between 3-6 days (Figure 2). This was the exception for one female whose vaginal membrane remained perforated for approximately 2 weeks after initially being observed as open. Despite this, she was still placed with a male as described below.

**Figure 1.**
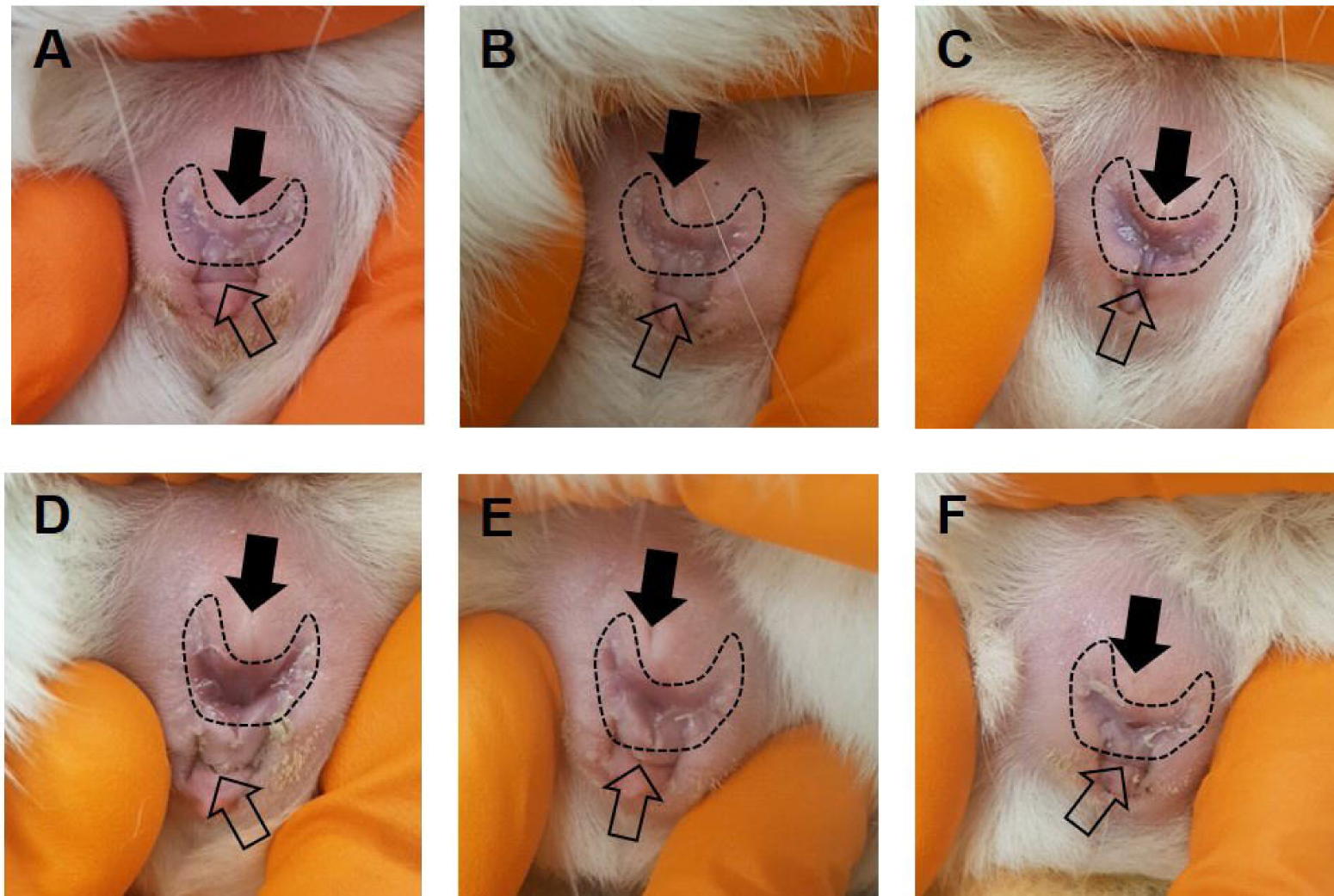
Monitoring of the guinea pig vaginal membrane. During the latent period of the estrus cycle, the guinea pig vaginal membrane (dashed outline) is visibly closed (**A** & **B**). 4-5 days prior to ovulation, changes to the color of the vaginal membrane can be observed indicating potential membrane perforation (**C**). At the time of ovulation, the vaginal membrane perforates and increased vaginal secretions/mucus can be observed (**D**). Following ovulation, the vaginal membrane begins to close (**E** & **F**). Closed arrow indicated urethral opening, open arrow indicates anus.

**Figure 2.**
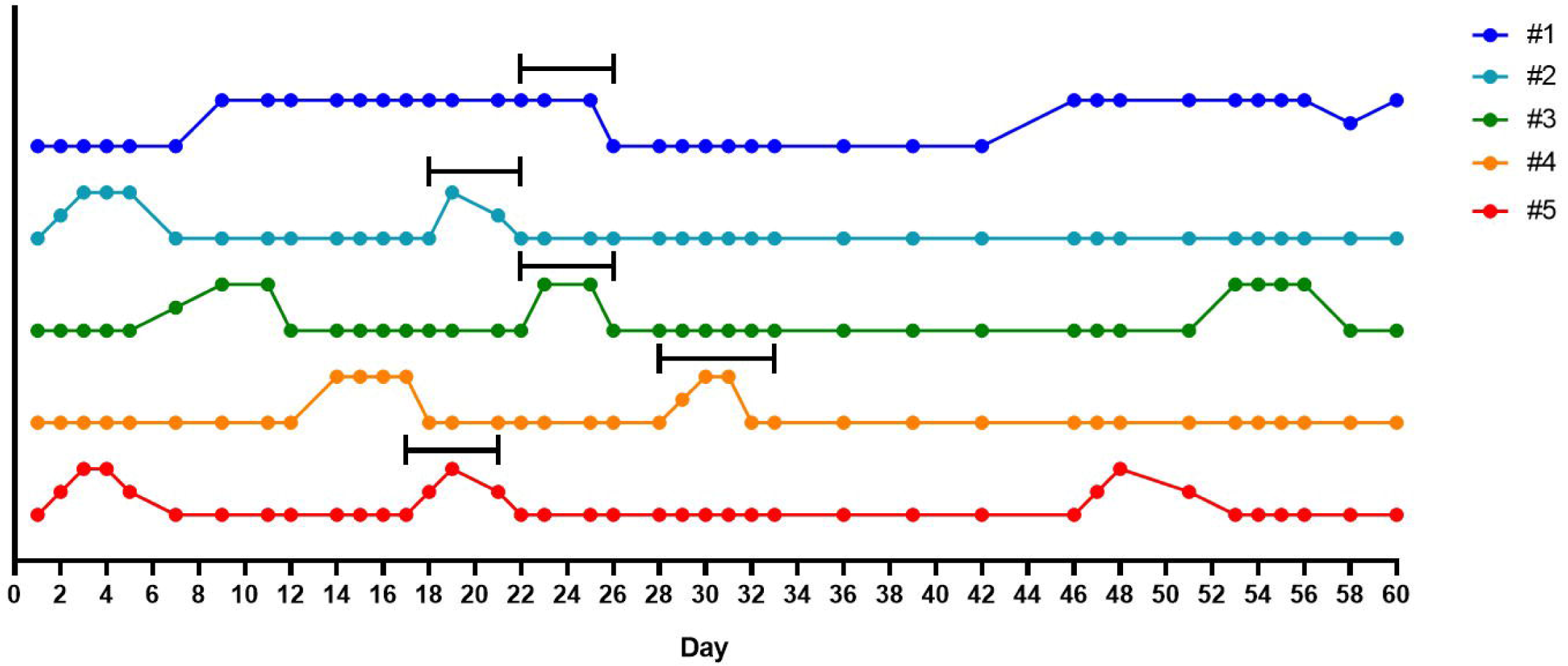
Graphical representation of the estrus cycle in the guinea pig based on monitoring of the vaginal membrane. In five animals, the vaginal membranes were monitored daily, a closed vaginal membrane was designated baseline and perforation of the membrane marked as an increase away from baseline. Females were placed with a male 14 days after perforation of the vaginal membrane was first observed as indicated by the back bars above the colored lines. Monitoring throughout pregnancy also occurred in which, the vaginal membrane was observed as perforated in 2 (#3 and #5) out of the 5 animals. For female #1, observations of a perforated vaginal membrane on day 46 also included blood in the vaginal lumen and a decrease in maternal weight gain. Later ultrasound confirmed loss of pregnancy in this animal.

### Time mating guinea pigs and determining guinea pig pregnancy

By monitoring changes to the vaginal membrane, we achieved a 100% pregnancy success rate with first time mating of 5 female guinea pigs. Females were placed with a male 14 days after full perforation of the vaginal membrane was noted. This was while the vaginal membrane remained closed but early changes, as previously described, were observed and allowed the breeding pair to become acquainted before copulation. During mating, daily vaginal monitoring continued and gestational day 1 was designated when the vaginal membrane was observed as fully perforated with what has previously been described as ‘estrus fluid’ [12]. The female remained with the male until the vaginal membrane closed; 5-6 days with the male total.

### Early pregnancy ultrasound to confirm pregnancy

Whilst it is relatively easy to observe the estrus cycle of the guinea pig by monitoring the vaginal membrane, determining a successful pregnancy is more difficult. Guinea pigs carry weight around their abdomens making visual observations of pregnancy practically impossible until after gestational day 40. Furthermore, noticeable increases in maternal weight, that would suggest a pregnancy, do not occur until after mid-pregnancy and can vary depending on litter size (Figure 3). Daily monitoring for vaginal membrane perforation is also not a viable technique as we have observed this phenomenon occurring in pregnant animals (Figure 2). In our experience, ultrasound is the only definitive method of determining pregnancy prior to gestational day 30 (Figure 4) but is limited to not being accurate before gestational day 21.

**Figure 3.**
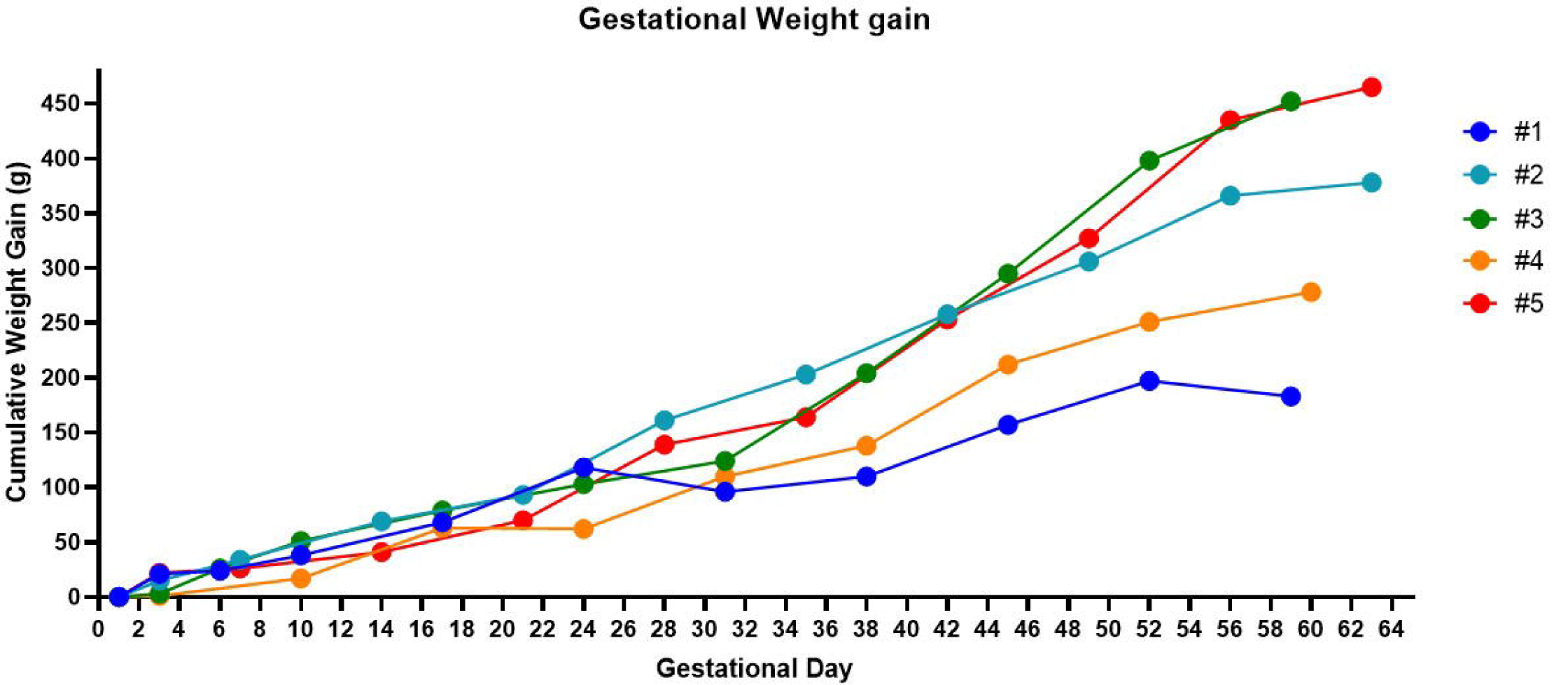
Weight changes in the guinea pig throughout gestation. Noticeable increases in gestational weight are not reliable until mid-pregnancy in the guinea pig. Healthy fetuses were confirmed in all animals at gestational day 30 using ultrasound, except for one animal (#1) in which blood in the vaginal lumen was observed at gestational day 24 and was associated with a decrease in cumulative weight gain. At post mortem, #3 had 4 fetuses, #5 had 3 fetuses, #2 and #4 had 2 fetuses and #1 was not pregnant.

**Figure 4.**
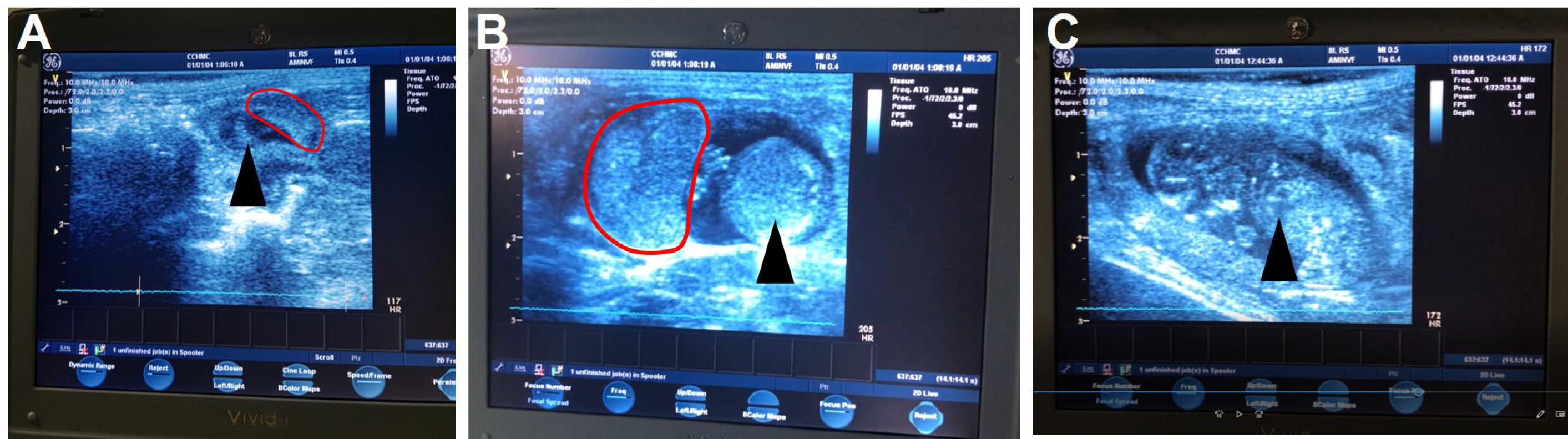
Ultrasound images of the guinea pig pregnancy. The earliest time to reliably detect a guinea pig pregnancy using ultrasound is approximately gestational day 21 in which the most obvious signs of a fetus are the amniotic cavity and placenta (**A**). By mid-gestation (gestational day 30-35) the placenta and sub-placental region are visible along with the fetus (**B**) including clear identification of the fetal heart (**C**). Black arrow identifies the developing fetus, pointing to the fetal heart in **C**. The red outline indicates the placenta.

## Discussion

The discovery and development of new treatments for obstetric complications is now at the forefront of reproductive medicine. As such, the requirement for appropriate pre-clinical animal models to evaluate the pregnancy response to treatment is imperative to ongoing research [4]. To date, mice and rat models commandeer most of the reproductive literature due to their ability to reproduce easily, reliably, and frequently. However, key aspects of reproduction in mice and rats; for example hormonal changes in the maternal environment, implantation and placental development, do not closely resemble human reproduction [18]. Furthermore, mice and rats are litter bearing species, and can carry 6-12 fetuses, as opposed to humans that generally carry singleton pregnancies. Therefore, particularly in studies with human translational potential, there is a necessity to use more comparable species like the guinea pig [4, 10].

In this technical report, we confirmed and demonstrate changes to the guinea pig vaginal membrane throughout the estrus cycle that others have described [12, 19, 20]. Furthermore, we show how this method of monitoring the guinea pig vaginal membrane can lead to successful timed-mating. Most interestingly, we found in 2 of the 5 females we tracked, perforation of the vaginal membrane during pregnancy occurred despite previous observational studies suggesting that closure of the vaginal membrane persists throughout pregnancy [12, 19]. This phenomenon has previously been described with the original hypothesis that opening of the membrane during pregnancy was associated with fetal loss [20]. However further studies concluded perforation of the membrane during pregnancy was a natural process unique to guinea pig and possibly driven by changes in estrogen and progesterone [16]. Our study supports this conclusion as early pregnancy ultrasounds showed no evidence of fetal resorptions in the animals whose vaginal membranes perforated during pregnancy. This was the exception in one female in which blood within the vaginal lumen was observed at gestational day 24 and was associated with a reduction in maternal weight gain. Guinea pigs do not exhibit pseudopregnancy following sterile mating [11] and as such, our observations with this animal indicated loss of the pregnancy which was later confirmed by ultrasound. Overall, the exact reason as to why vaginal membrane perforation occurs during pregnancy, and is restricted to certain individuals, is still unclear, and further adds to the difficulty in determining whether mating attempts result in successful pregnancies.

This study was focused on describing an easy method to time mate guinea pigs. However, as with all animal models, simply placing a female with a male does not always produce a pregnancy. One of the major advantage’s guinea pigs offer over other rodent species is increased gestational length and precocial development, ∼ 69 days vs post-natal developers of ∼ 21 days(mice and rats), however, more work is starting to be done using chinchillas which have a 105 days gestation. Particularly for translational research, and studies developing potential therapeutics, the increased length of gestation offers ample time to manipulate, treat and assess the interventional outcome [4]. However, with increased gestational length comes longer waiting periods, particularly when determining a successful mating. In mice, increased maternal weight due to pregnancy is only noticeable at mid-gestation [21] and is often used as a method of confirming pregnancy. In our study, a similar outcome was observed in pregnant guinea pigs with noticeable increases in maternal weight observed after gestational day 35. However, this is a significant amount of time to wait to establish whether re-mating is required. Some have described methods of gently palpating the maternal abdomen as early as gestational day 15 to feel for conceptuses [22] however, we have not been successful in using this methodology. Instead, we use early pregnancy ultrasound to confirm pregnancy. In our experience, gestational day 21 is the earliest time in pregnancy in which we can reliably visualize the developing embryo, but does provide an earlier indication of a successful pregnancy over other methods.

Integral to reproductive research is the ability to time mate animals in order to monitor and collect data at similar points throughout gestation. We report an easy and reliable method of time mating guinea pigs through daily monitoring of the vaginal membrane. A significant advantage of the method is that this mating strategy can be performed by a single researcher and does not require overly specialized tools or equipment and negates the need to perform daily, time-consuming smears in order to track estrus. Coupled with ultrasound as a secondary method to confirm pregnancy, we have shown that through familiarizing ourselves with the changes that occur to the vaginal membrane, we can achieve a 100% successful pregnancy rate in a matter of weeks, which ultimately reduces overall costs associated with animal husbandry and increases research success.

## Methods

### Ethics statement

Animal research approval was obtained from the Institutional Animal Care and Use Committee of Cincinnati Children’s Hospital Medical Center (Protocol number 2018-0065).

### Animal care, vaginal observations and mating

Female Hartley guinea pigs were purchased at 500-550 g (*Charles River*) and group housed on a 12 h:12 h light on/off cycle at 72°F, 50% humidity, with food and water provided ad libitum. Starting one week after arrival, females were weighed weekly and progression through the estrus cycle documented with daily monitoring of the vaginal membrane. These daily inspections occurred in the morning at approximately the same time each day. Females were placed with a male guinea pig (Hartley, approximately 12 months of age) 14 days after the vaginal membrane was first observed as perforated. During the period in which the female was with the male, daily vaginal monitoring continued to ensure perforation of the vagina. Ovulation was presumed to have occurred at the time in which ‘estrus fluid’ was observed, and any potential signs of mating such as a pale crustiness around the vaginal opening was documented. Females remained with the males until the vaginal membrane was observed to be closing or closed and then returned to the group housing.

### Confirmation of pregnancy using ultrasound

Pregnancy was confirmed by transabdominal ultrasound under anesthesia, but can be performed without anesthesia [23], 21-30 days after vaginal perforation was observed. General anesthesia was induced using 4-5% isoflurane mixed with 2 L/min oxygen and maintenance at 1 L/min and temperature and pulse oximetry monitored throughout the procedure. The female was placed on her back with the abdomen shaved. Ultrasound coupling gel was applied and pregnancy confirmation performed using a Voluson I portable ultrasound machine (*GE*) with a 25 E 12 MHz vascular probe (*GE*) on the cardiac setting. Following confirmation of pregnancy by visualizing a conceptus, the female was allowed to recover in an incubator before being returned to group housing. The whole procedure was performed in less than 20 min with the longest process being anesthesia induction.

## Declarations

### The authors declare no conflicts of interest

RLW designed and performed the protocol, analyzed the data and wrote the manuscript. KL performed the protocol and edited the manuscript. BJM and TRHR provided technical assistance and edited manuscript. HNJ provided monetary support, technical assistance, and edited the manuscript. All authors approved the final manuscript.

This study was funded by Eunice Kennedy Shriver National Institute of Child Health and Human Development (NICHD) award <u>R01HD090657</u> (HNJ).

